# Outside-in: intracellular vesicles in giant sulfur bacteria contain peptidoglycan

**DOI:** 10.1101/2022.04.11.487978

**Authors:** Beverly E. Flood, Dalton J. Leprich, Ryan C. Hunter, Nathalie Delherbe, Barbara MacGregor, Michael Van Nieuwenhze, Jake V. Bailey

## Abstract

Until recently, the cellular envelopes of bacteria were regarded as static and rigid relative to those of eukaryotes. While investigating peptidoglycan synthesis in populations of giant sulfur bacteria, *Candidatus* Thiomargarita spp., we observed internal vesicle-like features (VLFs). VLFs, as imaged following the active incorporation of D-amino acids, appear to begin as invaginations and delaminations of the cellular envelope. Staining with wheat germ agglutinin confirmed the presence of peptidoglycan in VLFs, while polymyxin B revealed that the outer membrane is present in some VLFs. Transmission electron microscopy revealed a complex network of interconnected VLFs. Genomes of *Ca*. Thiomargarita nelsonii lack a canonical divisome, while possessing homologs to genes such as actin, membrane scaffolding proteins, and dynamins that are associated with phagocytosis in eukaryotes. The physiological role of VLFs remains unclear, but the presence of sulfur globules in some suggests compartmentalization of metabolism and energy production. This is the first report of peptidoglycan and outer membrane bound intracellular vesicles within prokaryotic cells. These findings transform the canonical view of the inflexible bacterial cell envelope and further narrow the divide between prokaryotes and eukaryotes.

## Introduction

Members of the Beggiatoaceae are sulfur-oxidizing bacteria that are some of the largest bacteria in the world. Morphologically, these bacteria resemble cyanobacteria, a phylogenetically-distant group with which they have undergone substantial horizontal gene transfer [1–4]. *Ca.* Thiomargarita spp. include the largest known bacteria, with individual cells reaching up to a millimeter in diameter [5–8], (Fig 1). *Ca*. Thiomargarita cells appear hollow, with a single central vacuole occupying the majority of the cell volume [7, 8]. Like diverse marine eukaryotic phyla [9] and some sister marine Beggiatoaceae [10], *Ca*. Thiomargarita stores nitrate in a large central vacuole at high concentrations relative to the surrounding seawater [7, 11]. Nitrate serves as a terminal electron acceptor for the oxidation of sulfide in the absence of sufficient oxygen. The central vacuole may also store additional substrates and provide additional function(s) [12] but investigative studies are lacking. *Ca*. Thiomargarita spp. also have the capability of carrying out other types of lithotrophic and heterotrophic metabolism [7, 8, 13–16]. They store abundant inclusions of elemental sulfur, polyphosphate, and glycogen in the cytoplasm surrounding the vacuole, along with genetic material that is thought to include thousands of copies of a cell’s chromosome, reviewed by [8].

**Fig 1.**
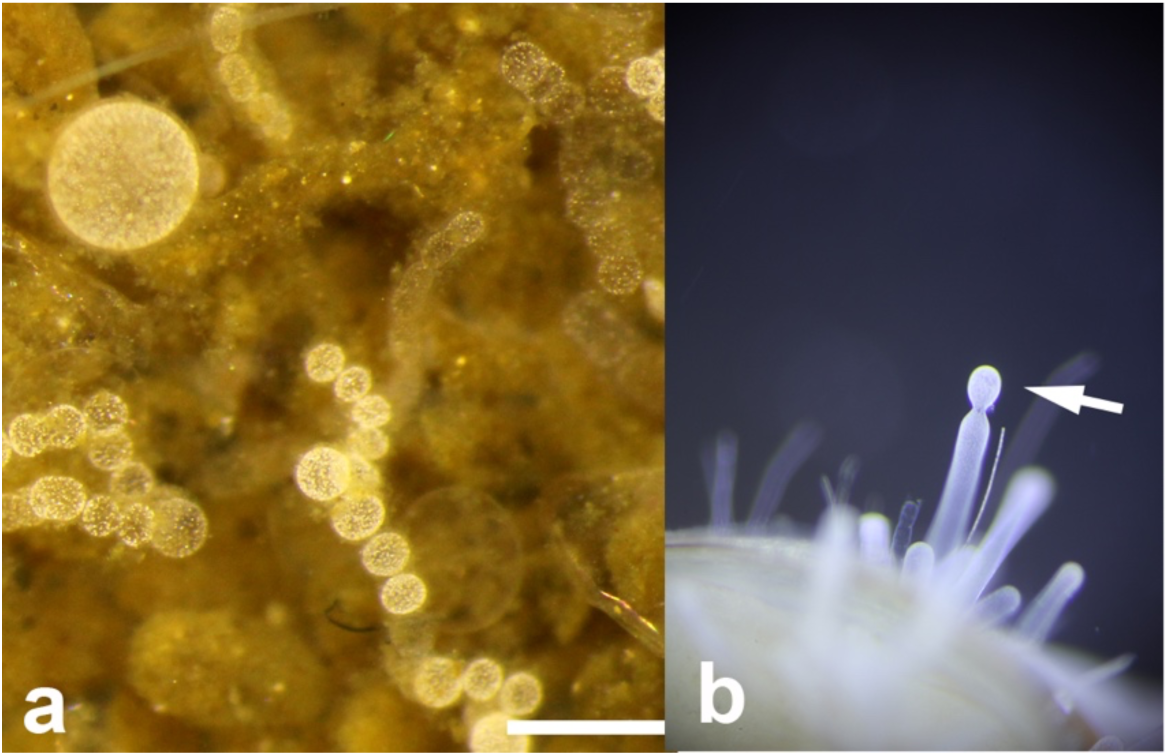
Examples of *Ca*. Thiomargarita spp. used in this study. (a) *Ca*. Thiomargarita spp. on collection along the Namibian margin exhibit abundant cells in various stages of division and diversity of cell shape and size. (b) *Ca*. Thiomargarita nelsonii attached to a Provannid gastropod collected from Hydrate Ridge, Pacific Ocean. A genome of this strain was included in genome analyses. Scale bar is 600 µm for both images.

Despite their macroscopic size, *Ca*. Thiomargarita have yet to be cultivated as isolates in the laboratory. But, some strains of *Ca*. Thiomargarita can remain viable in the laboratory for more than two years under refrigeration in their host sediments. Two genomes produced for *Ca*. T. nelsonii, as well as labeling experiments for oxidoreductase activity and microsensor measurements, suggest that these organisms have diverse metabolic potentials and are metabolically active following collection [11, 13–15]. Previously, a maintenance medium containing vitamins and trace metals along with lithotrophic or heterotrophic electron donors was shown to support metabolic activity in *Ca*. Thiomargarita spp. [11, 15]. Here, we incubated *Ca*. Thiomargarita with fluorescently-labeled D-amino acids (FDAAs) to visualize active peptidoglycan synthesis [17–19]. Peptidoglycan (PG), which provides mechanical strength to the cell ultrastructure, is composed of glycan strands linked by peptide chains containing D-alanine and D-glutamine [20, 21]. Fluorescently- labeled D-alanine [22] has been used in a variety of studies to facilitate the understanding of the formation and structure of PG, bacterial growth patterns, and how morphologically complex cells modulate their growth patterns (reviewed by [19]. Importantly, incorporation of labeled D-alanine is specific to PG, which is not labeled with the fluorescent enantiomer L-3-amino-L-alanine [18].

In this study, FDAA labeling was initially employed to better understand the rate of metabolic activity in *Ca*. Thiomargarita spp.. We hypothesized that due to their native habitat and the lack of observed growth *in vitro* [8], long incubation periods with FDAAs would be necessary because *Ca*. Thiomargarita are extremely slow growing. Furthermore, we hoped to observe how PG synthesis contributed to their morphology and modes of cell division (reviewed by [4]). Two previous *Ca*. T. nelsonii genomes [13, 14, 23] and two genomes produced for this study revealed that they possess the canonical genetic potential to synthesize PG, with the exception of a candidate gene for the key penicillin-binding protein 3 (*ftsW*). However, *Ca*. Thiomargarita lacks most of the canonical genetic repertoire for cell division, except a candidate gene for the septal ring tubulin homologue FtsZ. As such, we hypothesized that *Ca*. Thiomargarita spp. undergoes observable non-canonical cellular division.

## Results

### FDAA-labeling reveals active incorporation of PG along division planes

PG synthesis is required for cell wall elongation and the formation of the division septum. We observed incorporation of FDAAs into the *Ca*. Thiomargarita spp. ultrastructure as a weak but specific fluorescent signal along the cell margins, as well as an intense fluorescence signal localized at the division septum. FDAA incorporation at the division plane was observed in both cell trichomes (Fig 2b-c) and individual dividing cells (Fig 2a). Observable label was present in as little as fifteen minutes, suggesting very active metabolism (Figs 2b-c, S1 Movie). Label localization was similar to that observed in FDAA-stained bacteria in earlier studies, e.g. [17]. Labeling experiments conducted for one week revealed that dead cells do not incorporate FDAA. These observations show that FDAAs do not exhibit non-specific binding in dead cells, (S1 Fig).

**Fig 2.**
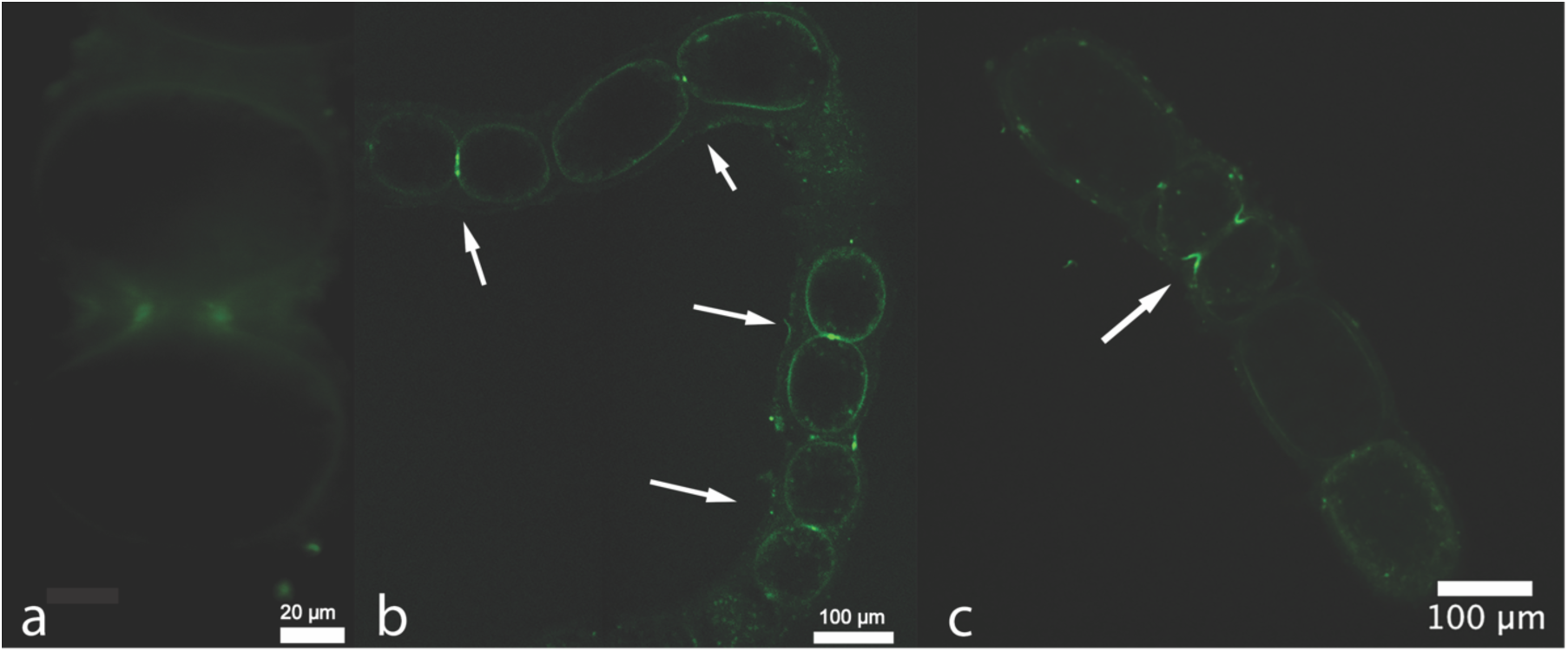
*Ca*. Thiomargarita spp. preferentially incorporate FDAAs along the division septum. (a) Cells incubated for 1 week under anoxic conditions with formate as an electron donor exhibited evidence of active cell division. (b) Confocal slice of cells exposed to FDAAs in the laboratory for 15 minutes, cells that appear to be dividing exhibit a preferential incorporation of FDAAs along the division septum (arrows) suggesting active PG synthesis associated with cell division. (c) Confocal slice of cells incubated with FDAAs for 15 minutes revealed reductive division.

### FDAA labeling reveals incorporation of peptidoglycan in VLFs originating from the cellular envelope

Active incorporation of FDAA into PG revealed that in some *Ca*. Thiomargarita cells, the cellular envelope budded inward to produce intracellular “pockets” (Figs 3 and 4). The cellular envelope surrounding the budding vesicles also exhibited active incorporation of the FDAAs (Figs 3a-b). In some cases, these pockets appeared to completely bud off from the cell envelope and became independent intracellular VLFs, (Fig 4a-c). Some unattached intracellular VLFs also exhibited active incorporation of FDAAs and were located either in the cytoplasm, or in the vacuole as observed in 3D reconstructions from confocal z-stacks (Fig 4d, S2 and S3 Movies).

**Fig 3.**
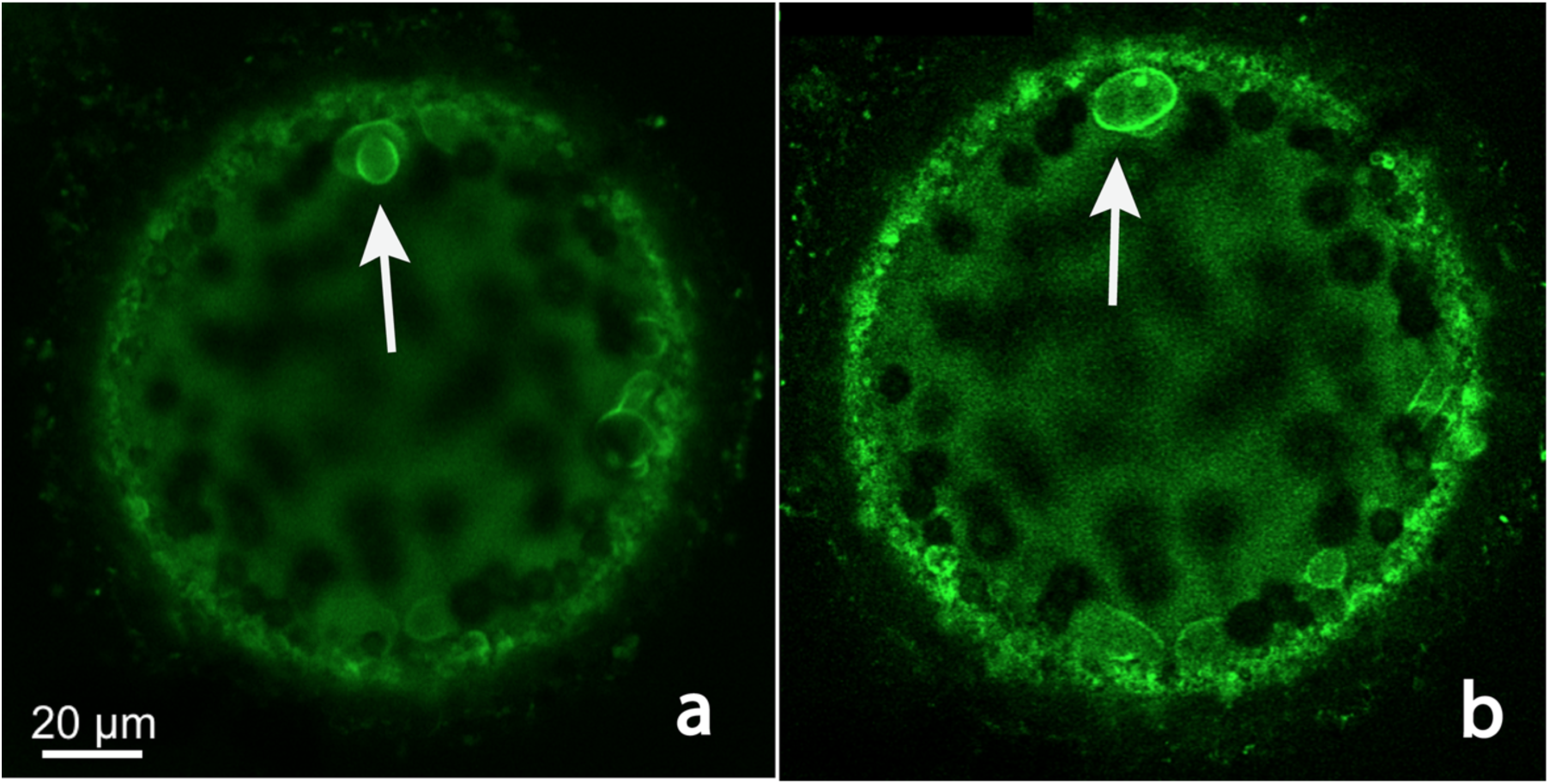
FDAA labeling of PG reveals intracellular vesicles form from the cellular envelope. The incubation time with the labels was 15-30 minutes. Dark intracellular spheres are sulfur granules. (a) A confocal microscopy slice through a *Ca*. Thiomargarita cell revealed the invagination of the cellular envelope producing an internal pocket/vesicle (white arrow) that has almost completely detached from the cell wall. (b) The same *Ca*. Thiomargarita cell as in (a) but a confocal microscopy slice closer to the surface of the cell revealed that there was active incorporation of FDAA in the cellular envelope surrounding the internal vesicle formation (white arrow). Note the base of the vesicle and a distal ring of PG around it exhibit the highest amount of incorporation of the FDAA.

**Fig 4.**
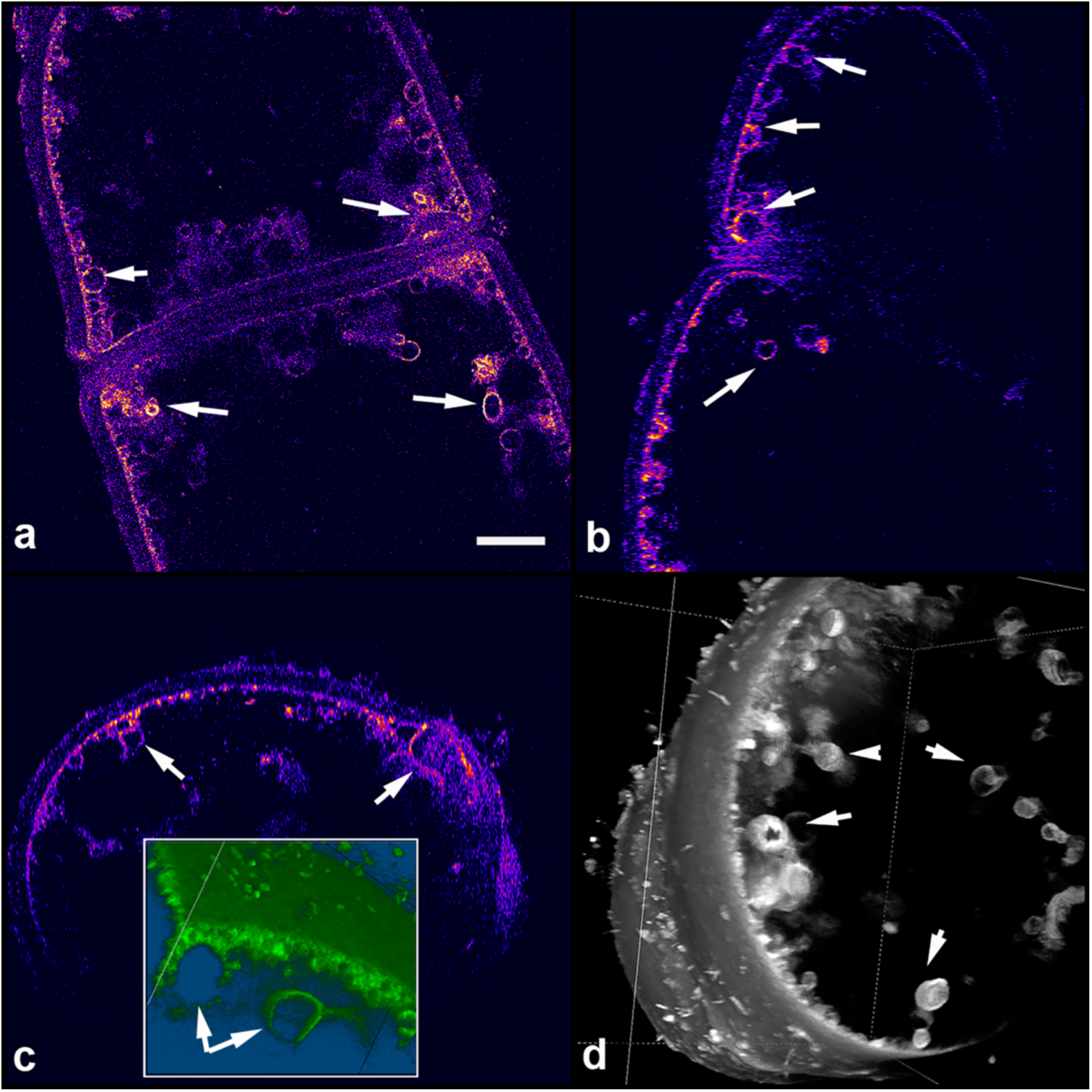
Vesicle-like features bounded by PG are actively maintained even after detaching from the cellular envelope. (a-c) Views along three axes of a confocal z-stack of FDAA-labeled *Ca*. Thiomargarita cells undergoing cell division. Invaginations and vesicles are indicated with white arrows. (d). A maximum intensity projection 3-D rendering of the cells in (a-c) showing the vesicles as spheres apparently translocated from the ultrastructure into the vacuole. Scale bar for all is 10 µm.

### Wheat germ agglutinin labeling for PG

*Ca.* T. nelsonii has the genetic potential to produce metabolites lacking L-stereo-specificity via non-ribosomal peptide synthetases. We considered the possibility that the vesicle-like

features are the products of D-amino acid incorporation by peptides synthesized outside the ribosome, or by other molecules not previously known to incorporate FDAAs. Wheat germ agglutinin (WGA) binds specifically to N-acetylglucosamine, a major structural unit of PG, so this test provides a chemically distinct line of evidence for PG bounding the VLF. Additionally, while FDAA labeling only reveals active PG synthesis, WGA staining provides visualization all PG within the cells. The results of the WGA staining were consistent with FDAA-labeling, and revealed some additional cellular features (Figs 5a-b). In some cases, we observed multiple layers within the cellular envelope that stained for N-acetylglucosamine and sometimes appeared to be delaminating. Additionally, sulfur globules appear to be bound within PG bearing vesicles and vesicular masses (Fig 5c and S4 Movie). Counterstaining with DAPI revealed that most DNA is located within a thin region near the cellular envelope, but some DNA is located surrounding large vesicles (Fig 6 and S5 Movie).

**Fig 5.**
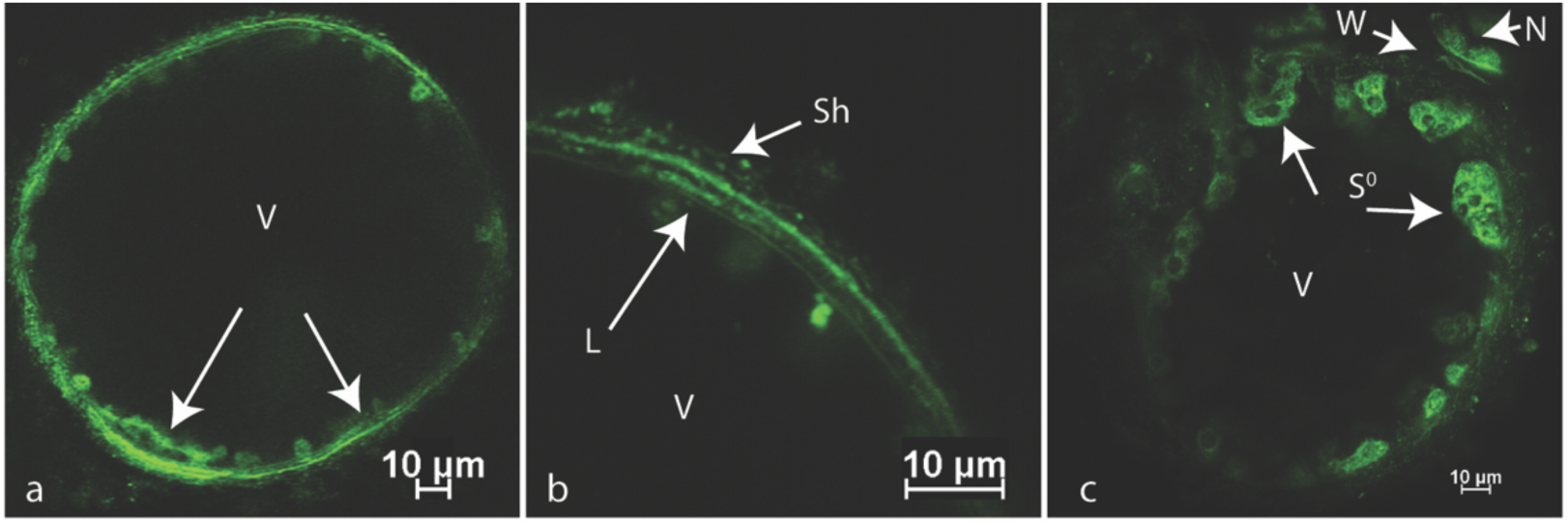
Whole cell staining with WGA revealed additional PG bearing features. V = vacuole. (a) Confocal microscopy slice through a *Ca*. Thiomargarita spp. cell revealed possible delamination of PG layer. (b) Increased magnification of cell in (a) revealed multiple layers and possible delamination, Sh = sheath with epibiont bacteria. (c) Confocal microscopy z-stack revealed PG bearing accumulating vesicles with some containing sulfur granules within them (dark masses within green), S0 = sulfur granules, W = PG wall, N = neighboring cell.

**Fig 6.**
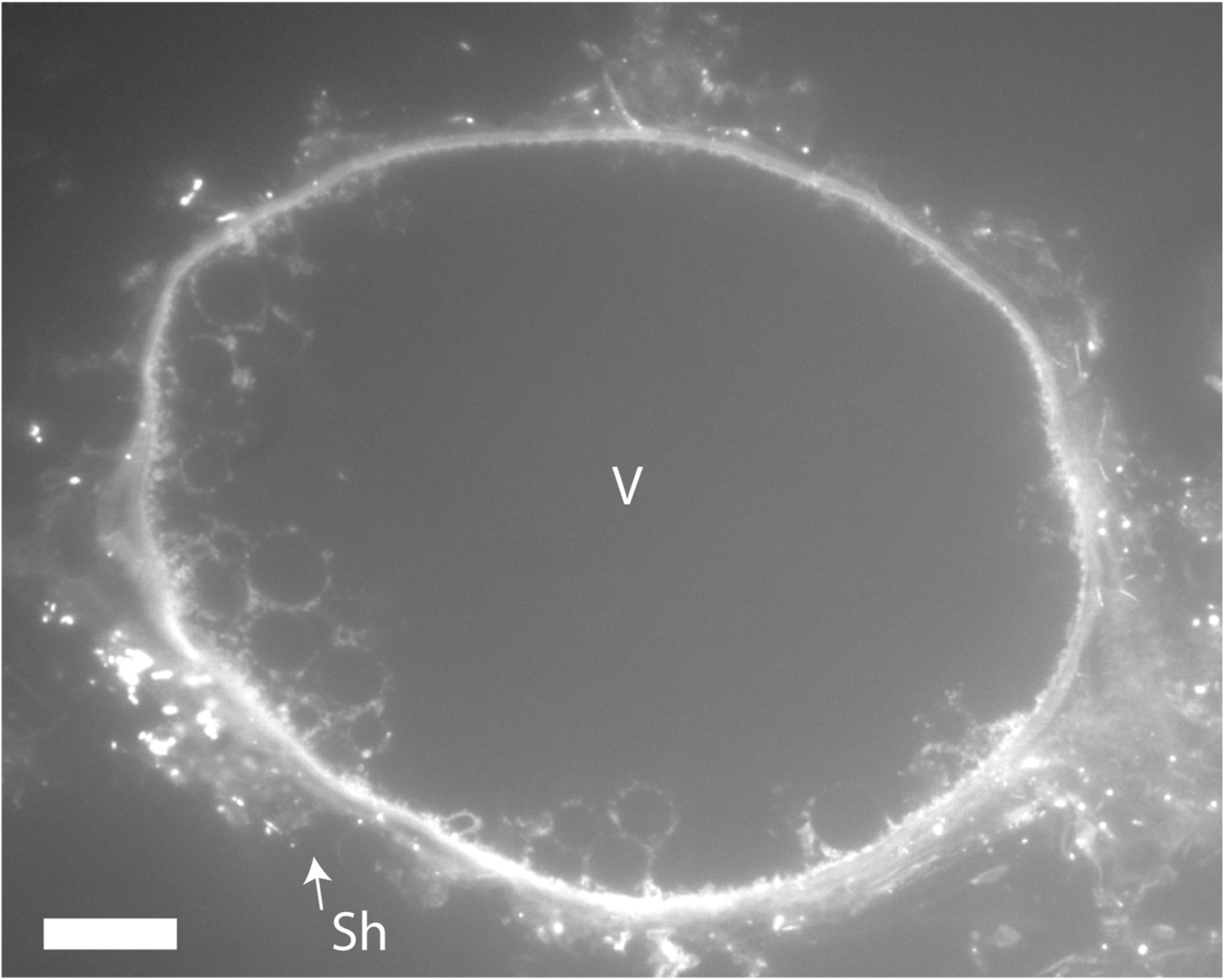
DAPI staining of thin sections reveal DNA predominantly in the cytoplasm but also associated with vesicles. V= vacuole, Sh = sheath with epibiont bacteria, scale bar = 20 µm.

### Fluorescent labeling of thin sections revealed other features of VLFs

Co-staining of thin sections with polymyxin B, which binds specifically to lipid A in the outer membrane, and FM 4-64, a general lipid counterstain, revealed that outer membrane lipids are also present in some but not all intracellular vesicles (Figs 7a-b). The optical resolution of light microscopy was insufficient to confirm that outer membrane lipids were inside the vesicles, but the fluorescence of polymyxin B did encroach further into the vesicles than that of WGA, which supports the possibility that they are inside as would be predicted if the entire ultrastructure inverted during infolding.

**Fig 7.**
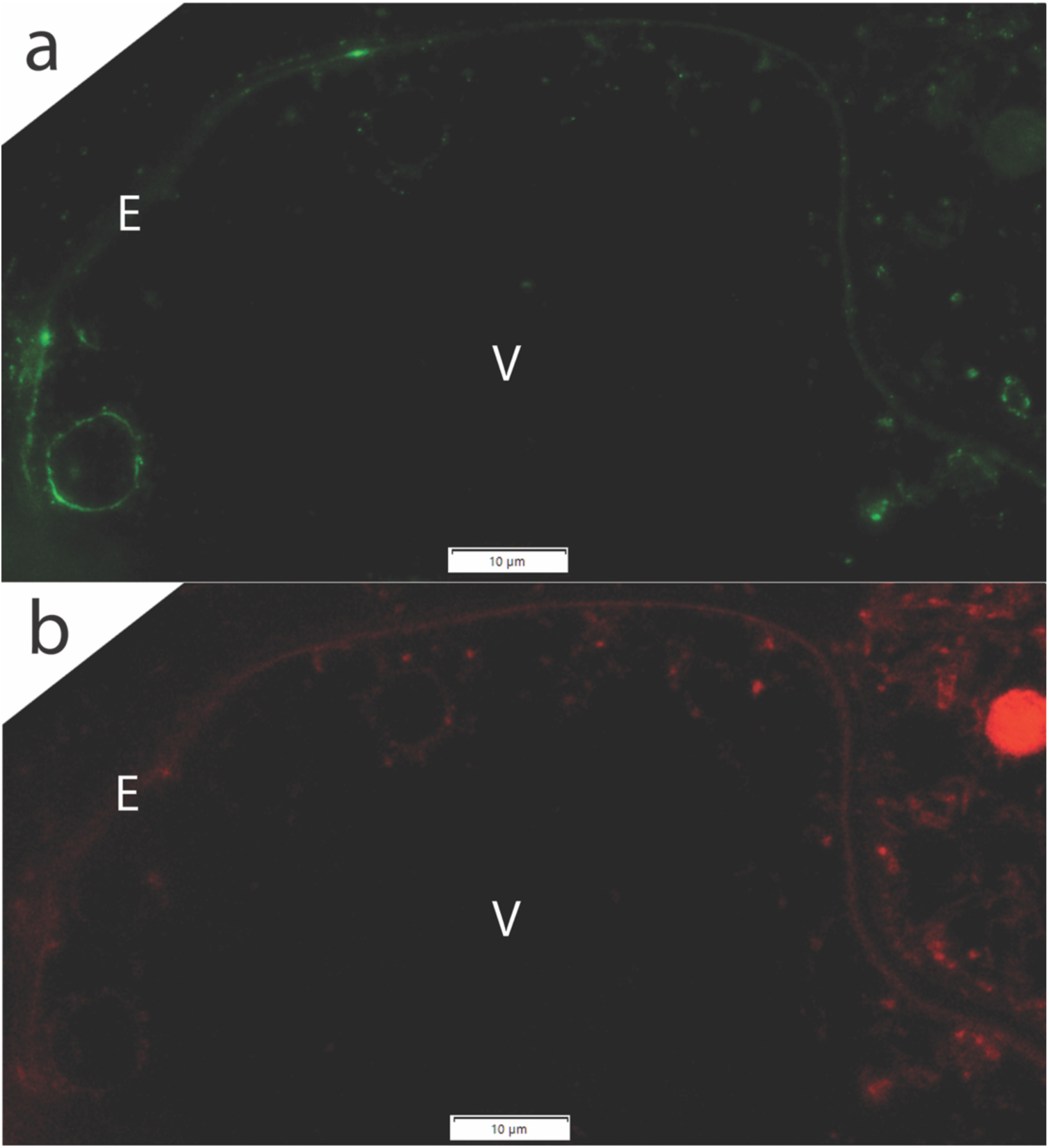
Thin sections of *Ca*. Thiomargarita spp. reveal that vesicles sometimes include outer membrane material. (a) lipid A stained with polymyxin B (b) lipids as stained with FM4-64, E = envelope, V = vacuole.

### Transmission electron microscopy

Two distinct fixation and staining methods were employed to address challenges with preserving the cellular structure of such large cells with vast aqueous interiors that are prone to collapse e.g. (S2 Fig). TEM revealed VLFs, as observed with light microscopy, as well as additional features (Fig 8 and S2 Fig). The VLFs varied in size from sub-micron to more than 50 µm in diameter. While sometimes the VLFs contained sulfur, often they contained other storage products such as electron-dense inclusion bodies, DNA, and ribosomes, but most appeared devoid of material that stained with contrast agents. In general, the cytoplasmic region containing the VLFs appeared as a highly disordered network of interconnected vesicles of varying sizes. Some of these “networks” were detached from the main cytoplasmic region. The cellular envelopes, which were approximately 100 nm thick, consisted of at least five layers. The darkest and thickest stain layer, presumably PG, was the second observed layer from the interior. The PG layer was 6-10 nm in thickness, which is thicker than some bacteria [24]. Occasionally, VLF envelopes were contiguous with the cellular envelope or appeared contiguous with envelope delaminations. The thickness and electron density of VLFs were highly variable. Some VLFs envelopes appeared somewhat disordered and spongy, while other envelopes were well-defined layers. A second, thinner layer was sometimes observed interior to the PG-like layer of the VLF. Lastly, we also observed extracellular bilayer vesicles which were distinctly different in morphology from the VLFs, (S3 Fig).

**Fig 8.**
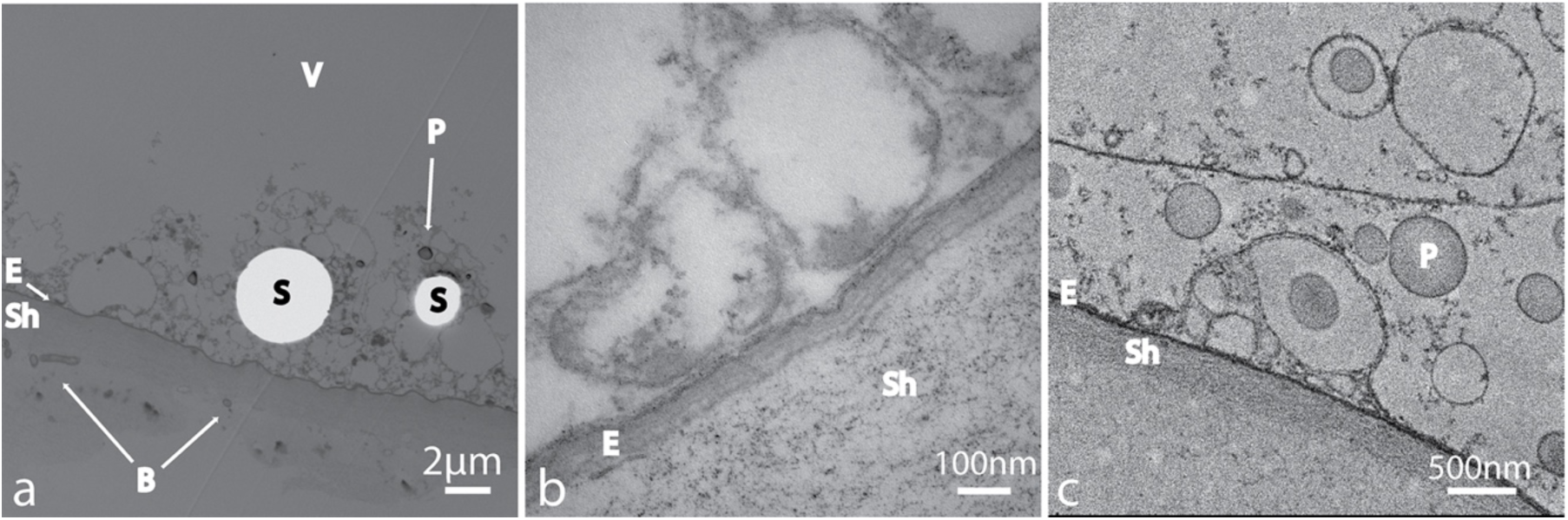
Additional evidence for PG in vesicle-like features was provided by TEM. (a) TEM section demonstration heterogenicity in vesicles. Some vesicles contain sulfur granules (S) but most do not. (b) The cellular envelope (E) is complex consisting of five or more layers and is almost 100 nm thick. (c) The vesicles envelopes are sometimes on contiguous with the cellular envelope and can consist of two or more layers. Here vesicles are sometimes surrounding polyphosphate granules (P) but not always. B = epibiont bacteria, E = envelope, P = polyphosphate Sh = sheath, and V = vacuolar region. Images (a) and (b) were samples processed via TEM method #2 and (c) method #1.

### Genome analyses

Four *Ca*. T. nelsonii genome bins, two of which were produced herein, were examined for the genetic potential to synthesize outer membrane lipopolysaccharides, a tethered outer membrane (Tol-Pal System) [25, 26], meso-diaminopimelate-containing PG, a division septum, and to carry out cellular elongation [27]. But all genomes lack nearly all canonical genes associated with Z-ring formation, including those for septal PG synthesis (*ftsIW*) and proteins anchoring FtsZ to the cellular envelope (*ftsABEKLNQX*, *zipA*) (S1 Data). Given our unusual evidence for PG and OM present in some intracellular vesicles, we expanded our search to other potential cytoskeletal genes found in morphologically complex bacteria and their known associated genes, as well as with membrane remodeling and vesicle formation in eukaryotes. A complete account of genes queried are presented in S2 Data, with select results potentially related to VLF formation discussed below.

## Discussion

This project was initiated to investigate whether *Ca*. Thiomargarita spp. were metabolically active and undergoing cell division *in vitro*. Since we have not witnessed a single dividing cell exhibit notable changes in cell division in any of our lab’s periodic attempts to cultivate *Ca*. Thiomargarita spp. from our sampling sites off the coast of Namibia over the past 10 years, our initial hypothesis was that we would not see incorporation of FDAAs in short incubation periods as seen in classic laboratory strains [17, 18]. However, we detected incorporation of the FDAAs in as little as ten minutes, including localization along the septal plane. Thus, we conclude that *Ca*. Thiomargarita spp. are actively growing and dividing within refrigerated samples of their host sediments. Although artificial enrichment and cultivation attempts tested thus far have been inhibitory to complete cell division and long-term viability, our previous work using redox-sensitive dyes indicate that the cells are metabolically active for months to years following collection [15]. We also recently discovered that *Ca*. Thiomargarita hosts a large number of host-specific epibionts [28]. Consideration of these microorganisms, in addition to *Ca*. Thiomargarita-specific metabolic attributes, may be important in successful cultivation of *Ca*. Thiomargarita.

We were surprised to find that the labeling experiments revealed internal vesicle formation that included PG and sometimes the outer membrane. The co-occurrence of FDAAs in an attached vesicle and the neighboring cellular envelope (Fig. 2a-b) suggests that perhaps structural remodeling of the cellular envelope and/or overproduction of constituents of the envelope promotes vesicle formation. Internal compartmentalization derived from the cellular envelope in prokaryotes is not uncommon. But to our knowledge, no other intracellular bacterial vesicles have been shown to include PG or outer membrane constituents [29–31]; this includes several studies utilizing FDAAs on morphologically complex bacteria and those which produce intracellular vesicles, e.g. [32–40].

Comparison of numerous TEM studies on the Beggiatoaceae [8, 41–51] suggests that both the cellular envelope and the intracellular environment of this clade is highly heterogeneous, even between strains within the same genus. Some strains, including *Ca*. Thiomargarita spp., have up to five layers in the cellular envelope, suggesting the presence of S-layers. Marine Beggiatoaceae strains tend to exhibit a spongy intracellular appearance surrounding a central aqueous vacuolar region that is devoid of structure. In general, many of the Beggiatoaceae appear to have an extremely disordered intracellular environment where membranes, vesicles, and electron-dense materials like DNA cluster in what appears to be a disordered fashion. Within the same cell, a VLF may surround carbon stores or polyphosphate stores, but in other locations in the same cell, storage granules are found outside of VLFs, and VLFs that possess no storage granules are common. In the Beggiatoaceae, sulfur globules are usually found within membrane-bound vesicles thought to be derived from the cytoplasmic membrane. Interestingly, sulfur granule envelopes in *Beggiatoa alba* B15LD have been observed to be composed of up to five layers [44]. In *B. alba* B15LD the sulfur globule “envelope” was destroyed by lysozyme, which acts on PG [52]. But in most cases, multiple layers of ultrastructure were not observed in Beggiatoaceae sulfur globules envelopes or the surrounding membrane. The preparation for TEM can cause artifacts or loss of information in electron micrographs [53, 54]. Some researchers reported difficulty in preventing mechanical distortion of cells during TEM and in some cases, the problem of a “lack of differentiating materials” [41]. Others found that the method chosen for preparation of the sample resulted in the loss of information [43]. Indeed, we encountered all these issues with TEM preparation of Ca. *Thiomargarita* spp. and preserving intact cells using typical preparation approaches was a challenge, which is why we turned to microwave impregnation with low viscosity resin.

### Potential mechanisms for VLF formation

It is possible that increased levels of microdomain lipid synthesis together with alterations in or detachment from the PG sacculus could alone drive vesicle formation, as for outer-membrane derived extracellular vesicles (reviewed by [55, 56]). Indeed, L-form bacteria (a.k.a. cell wall-less bacteria) form both external and internal vesicles [57] and their replication is independent of an FtsZ-based septum [58]. Outer-membrane vesicles have been shown to contain PG, so the association of PG with vesicles is not without precedent [59]. PG was, until recently, thought to be a highly ordered crystalline substance. However, recent atomic force microscopy imaging shows that different bacterial morphotypes can have differing degrees of ordering to the cell wall, including differences in the length and orientation of PG [60]. Short glycan chain lengths may increase the flexibility of the PG sacculus [24, 55], which may permit folding of the cellular envelope.

Cardiolipin, a non-bilayer anionic phospholipid, generates a large curvature and tends be concentrated at the division septum and poles of bacteria, and is also enriched in the highly-invaginated inner membrane of mitochondria [61, 62]. Cardiolipin is thought to be inserted into the inner membrane leaflet, where it can bind to proteins. Cardiolipin has roles in controlling membrane fluidity, lipid bilayer remodeling, and deformation. But cardiolipin is ubiquitous in bacteria and VLFs are rare.

Recent work has shown that outer membrane vesiculation can be mediated by curvature-inducing proteins [63]. While periplasmic turgor pressure can provide for outer membrane blebbing, as in Fig. S3, it seems unlikely that an external force would supply the mechanical force needed to induce envelope infolding to form VLFs. Vesiculation in the formation of chromatophores of purple bacterium *Rhodobacter sphaeroides* is thought to be driven by CM lipid synthesis and remodeling, but these vesicles do not contain PG [64]. It is hard to imagine that lipid enrichment alone could be responsible for making large symmetrical PG-containing vesicles capable of completely detaching from the envelope.

### *Ca*. T. nelsonii has the genetic potential for endocytosis-like VLF formation

Eukaryotic endocytosis is not a passive process whereby lipid dynamics alone drive vesicle formation, reviewed by [65–67]. These mechanisms in eukaryotes are far too diverse and often not sufficiently-well understood to fully discuss herein, but key steps include: 1) *localized changes in membrane fluidity and tension involving both lipids and membrane scaffolding proteins*; 2) *actin or dynamins pinching off large vesicles from the envelope*; 3) *regulation of these processes by small GTPases*. Importantly, homologues of genes involved with endocytosis are known to occur in bacteria, reviewed by [68].

The dynamics of alterations of the membrane tension and fluidity in clathrin-independent endocytosis are not well understood, particularly the importance of different drivers. In general, membrane fluidity can be increased by packing the membrane with lipids such as phosphatidylserine, which increases the charge density. In *Ca*. T. nelsonii this may involve varying ratios of the long-tail fatty acids, phospholipids, and cardiolipin. To elucidate the role of membrane lipids in vesicle formation, an examination of the KEGG maps [69] for the biosynthesis of glycerophospholipids revealed *Ca*. T. nelsonii lacks the genetic potential to make choline and inositol phospholipids. Instead, they may make phosphatidylserines, phosphatidylethanolamines, and phosphatidylglycerols (although the final enzymatic phosphatase is missing, other genes of similar function can occur, e.g. [70]).

In eukaryotes, membrane tension resulting from lipid rafting is mediated by cholesterol and sphingolipids, and perhaps scaffolding proteins such as flotillin [71]. Lipid rafting was until recently thought to be restricted to the eukaryotes, but more recently it has also been demonstrated in model bacterial strains, reviewed by [72–74]. Bacterial lipid rafting may be mediated by sterols or other isoprenoids such as hopanoids and carotenoids, cardiolipin, and bacterial homologues to flotillin. The steroid and terpenoid biosynthesis KEGG maps revealed the genetic potential to synthesize cardiolipin, squalene, terpenoids, and perhaps sterols. *Ca.* T. nelsonii possesses two *orfs* encoding squalene synthases, two sterol desaturases/sphingolipid hydroxylases, and multiple *orfs* encoding putative sterols. Indeed, not all sterol synthesis genes are known [75] and new or alternative ones are regularly being discovered, e.g.[76]). The sterol identified in sister marine vacuolated taxon *Ca*. Marithioploca was cyclolaudenol, a C31 sterol which is found in a few plants and has not been reported in other bacteria nor in the marine environment [77]. We also found 1 flotillin and 13-14 flotillin-like *orfs*, including bacterial membrane scaffolding genes *hicCKX* [78]. The role of flotillins in endocytosis is currently under debate [65, 66]. In bacteria, flotillins have been shown to be in microdomains where they play a role in protein secretion, signal transduction, and transport. In *Bacillus subtilis*, the flotillin-like protein YdjI has recently been shown to be key to localization of the inner-membrane remodeler phage shock protein A, for which there are two *orfs* in *Ca*. T. nelsonii genomes [79].

There are many pathways for endocytosis that rely on the activities of actin, reviewed by [66]. For the sake of brevity, we focused our attention on the two mechanisms that can generate vesicles greater than 0.5 µm in size in eukaryotes, namely macropinocytosis and phagocytosis. In macropinocytosis, actin is responsible for making the vesicular cup via ruffling the plasma membrane [80]. Initially, a circular region rich in less-viscous lipids and containing a small signaling GTPase forms in the plasma membrane. Actin then polymerizes around the patch of fluid membrane, forming a ring capable of constriction. Macropinocytosis is unique among the types of actin-driven endocytosis because it does not always require dynamin to pinch off the vesicle, whereas phagocytosis does [81]. Another major difference between the two processes is that initiation of macropinocytosis is typically mediated by growth factors, whereas phagocytosis is mediated by external particle recognition. In both cases, aspects of formation, maturation and regulation of these envelope-derived intracellular vesicles are not fully understood.

A bacterial actin, MamK, has been implicated in intracellular vesicle formation to form magnetosomes in magnetotactic bacteria [82, 83] and a *mamK*-like *orf* has been previously identified in other Beggiatoaceae [23]. We found a *mamK* homologue in three of the *Ca*. Thiomargarita genomes. In some *Ca*. T. nelsonii genomes, the actin homologue *mamK* directly neighbors a flotillin-like homologue (pfam01145 Band 7). We also identified many *orfs* that contain uncharacterized heat shock protein 70 (hsp70) domains that belong to the actin superfamily. Hsp70 chaperone proteins are known to insert in negatively charged lipid-containing membranes (e.g. cardiolipin) and play a role in endocytosis among many other functions [84, 85], including membrane remodeling [86–88]. The hsp70 domain-containing *orfs* were particularly prevalent in the genome of the Ca. T. nelsonii strain from methane seeps that attaches to substrates, elongates, and buds new cells at the terminal end [5, 14]. With the exception of *dnaK* and *hscAC*, other actin homologues (e.g. *parM* [89]) were not found.

Dynamins are large GTPases that assemble into helical polymers that wrap around membrane tubes and contract upon GTPase activity. A bacterial dynamin was recently found to mediate membrane remodeling *in vitro* [90, 91], including membrane fusion. Dynamins have also been shown to stabilize FtsZ [92] and in some cases, also interact with a flotillin [93]. We found up to three dynamin-family protein encoding *orfs* in the *Ca*. T. nelsonii genomes. Thus, homologues of the key agents for completing vesicle formation in eukaryotic phagocytosis, actin and dynamin, are found in *Ca*. T. nelsonii genomes.

In both macropinocytosis and phagocytosis, a small Ras superfamily GTPase, pfam00071, regulates the actin. A Ras domain gene (pfam00071) occurs in all *Ca*. T. nelsonii genomes, where it resides on a possible operon containing a gene for ribosomal large subunit pseudouridine synthase B, a putative outer membrane/PG-binding encoding gene, and an *orf* of unknown function. In *Ca*. T. nelsonii Bud S10, from a Hydrate Ridge methane seep [14], this potential operon is adjacent to predicted genes for PG and *ftsZ.* Genes coding for proteins assigned to the Ras superfamily GTPase (pfam00071) appear to be uncommon in bacteria (361 of >7800 species in the pfam database) and archaea (38 of >300 species), but common in eukaryotes (989 of >1500 species). We also identified another small GTPase, pfam08477 and pfam16095, tentatively identified as “Roco” or “Rup” group ATPases [94]. This small GTPase is also uncommon in bacteria (177 species) and archaea (1 species) and is thought to be homologous to eukaryotic Ras proteins. In addition, we identified other potential small GTPases that could potentially function in cytoskeletal remodeling.

### *Ca*. T. nelsonii also has bacterial homologues to other eukaryotic cytoskeletal proteins

The genomes of *Ca*. T. nelsonii contain *orfs* encoding homologues for the cytoskeletal proteins tubulin [89, 95, 96] and intermediate filaments (IFs). Homologues of *α*-tubulin and *β*-tubulin were previously identified in sister taxa *Beggiatoa* sp. SS [96], but were not found in *Ca*. T. nelsonii. Instead, the tubulin homologue was of the ftsZI1 family (pfam13809), which has been hypothesized to be involved with membrane remodeling [95]. Intermediate filaments (IF) differ from actins and tubulins in that they are more deformable and elastic [97]. They self-assemble and do not bind a nucleoside triphosphate.

The structural composition of IFs is highly variable, but a coil-coil domain is essential for monomers to assemble laterally. Homologues to the heteropolymer-forming IFs ZicK and ZacK, recently discovered in the cyanobacterium *Anabaena* spp. [98], are also present in the *Ca*. T. nelsonii genomes. In *Anabaena* spp., ZicK and ZacK interact with MreB and cyanobacterial proteins SepJ [33, 99] and SepI [34], which are not found in *Ca*. T. nelsonii genomes. In *Anabaena* sp. PCC 7120 the heteropolymers span the length of the cell, longitudinal to the division septum, and are likely anchored to the poles. They contribute cell shape but not division. The model strain for colorless sulfur bacteria, *Allochromatium vinosum*, possesses an IF-like protein associated with sulfur globules, SgpG [100]. Homologues were not detected in *Ca*. T. nelsonii’s genomes nor did we find homologues to other known bacterial IFs [101–115]. The homopolymer bactofilin, CcmA (pfam04519), an alternative to IFs in bacteria [116], has been found broadly across bacteria and is present in *Ca*. T. nelsonii genomes. We speculate that tubulin or IFs or both may play some role in VLF formation.

### Is intracellular PG-bound vesicle formation unique to the *Ca*. Thiomargarita spp. and possibly the Beggiatoaceae?

Recently, a strain of Planctomycetes, *Ca*. Uab amorphum, was found to be capable of both locomotion and phagocytosis similar to that of some amoeba [117]. The authors attributed the behaviors to a eukaryotic or archaeal-like actin and demonstrated actin-like fibrous regions within the cell via TEM. We have not observed such structures in *Ca*. Thiomargarita spp. during our TEM investigations even though *Ca*. T. nelsonii possess an actin homologue. But presumably, phagocytosis by this strain of Planctomycetes involved deformation of the PG sacculus.

The surgeonfish gut symbiont, *Epulopiscium spp*., can approach the size of *Ca.* Thiomargarita spp.. A member of the Clostridales, *Epulopiscium* is unusual in that its envelope lacks a thick PG layer and it can produce multiple endospores. Furthermore, some cells have extensive and broken invaginations of the inner membrane that contain numerous nucleoids [118, 119]. This dense network of fibrous and membranous materials may function similar to a cell wall in providing structure [118] and may facilitate transport, overcoming limitations of diffusion for a large cell [120]. The function and molecular composition of this network is currently unknown [121]. The largest morphotype of *Epulopiscium* also possess a zone of “coated” vesicles hypothesized to be similar to eukaryotic clathrin-coated vesicles and excretionary in nature [118]. Unfortunately, the mechanism(s) for forming these intracellular vesicles is not known, nor are there suitable genomes to interrogate. The electron-dense VLFs in *Ca.* Thiomargarita spp. have a roughly similar appearance to vesicles observed within the cortex of the *Epulopiscium*, except that the VLFs in *Ca.* Thiomargarita spp. are primarily spherical, whereas *Epulopiscium* contains a layer of small spherical vesicles superjacent to a zone of irregularly-shaped vesicle-like features with electron dense boundaries [118]. The similarity in appearance between the VLFs in *Ca.* Thiomargarita spp. and *Epulopiscium* suggests the possibility that vesicles in *Epulopiscium* may also be bounded by PG and that PG-bounded VLFs may be a common feature of giant bacteria.

## Conclusion

This report represents an accumulation of data collected over of six years on *Ca*. Thiomargarita spp. from Namibian upwelling sediments. Their large size may be facilitated by the lack of the canonical cell division pathway. More importantly, we have learned that these extraordinarily large bacteria have the potential to bring the outside milieu inside their cells, as in eukaryotic endocytosis. However, the thick sheath and S-layer(s) of the envelope likely prevent most large particles and microorganisms from being drawn into the VLFs. We posit that one potential role of VLFs may be related to sequestering sulfur and other storage granules from the cytoplasm and from other vacuolar contents, although most vesicles do not contain inclusions. In particular, redox intermediates of sulfur oxidation, such as thiols [122], can react with reactive species generated by denitrification [123], which are found at high concentration in the *Ca*. T. central vacuole. Furthermore, in a sister taxon to *Ca*. Thiomargarita spp., *Ca.* Allobeggiatoa halophila, the central vacuole contains active electron transport chains that maintain the acidity of the vacuole by proton motive force while conserving cellular energy [124]. Thus, the VLFs in *Ca*. Thiomargarita spp., may function like cells within a cell, where cellular regulation and activity is localized and depends on the VLF’s internal content. DAPI staining indicates that DNA is associated with the vesicles, suggesting the possibility of locally-regulated gene expression. Such “nodes” of electron transport chain-mediated phosphorylation may prove beneficial in powering such a large cell. A third potential benefit to forming VLFs is to increase “surface to volume” ratios such that there is an increased number of energy-producing reaction centers to fuel the cell, as for thylakoids and chromatophores. Finally, another possible function of the VLFs may be to recycle outer membrane constituents, which would not otherwise be possible. Regardless of their metabolic function(s), the dynamic remodeling of the cell ultrastructure required to produce these features extends the boundaries of typical bacterial cell wall plasticity, adding to the growing list of unique features possessed by the largest of all bacteria.

## Methods

### Sample collection

*Ca.* Thiomargarita spp. were collected using a multi-corer on board the R/V *Mirabilis.* Cores were taken from organic-rich sediments on the Atlantic shelf, near Walvis Bay, Namibia (23°00.009’ 14°04.117’) on yearly expeditions from 2016-2019. All cells used in the experiments were collected from a depth of 1-3 cm depth beneath the sediment/water interface. *Ca*. Thiomargarita sp. cells were stored in their host sediments with overlying core-top water in closed 50 ml centrifuge tubes at 4°C and protected from direct light in advance of labeling experiments and TEM observations. For experimentation, individual chains of *Ca.* Thiomargarita spp. are separated from their host sediment in sterilized Instant Ocean*®* in a 35 mm sterile petri dish via pipet while observing them on an Olympus SZX- 16 stereomicroscope.

### Peptidoglycan labeling

*Ca.* Thiomargarita cells were incubated in the incubation medium described in [15] or in sterilized Instant Ocean*®*, along with 1 mM fluorescently-labeled amino acids and 1% dimethyl sulfoxide to improve the solubilization of the FDAAs [17]. The FDAA used in these experiments was BADA (4,4-Difluoro-5,7-Dimethyl-4-Bora-3a,4a-Diaza-s-Indacene-3-Propionic Acid-3-amino-D-alanine) EX 503 nm; EM 512 nm) Color Green/FITC [17, 22]. The cells were incubated in the FDAA media for 15 minutes to 1 hour in a sterile glass bottom petri dish prior to imaging. The cells in figure 1B-C were imaged using an Olympus IX-81 inverted microscope equipped with a long working distance (WD 2.7-4.0) 40x objective (NA-0.6), a DP73 camera and CellSens Dimension (Olympus, Japan). Other light microscope stacks were collected using a Nikon TiE inverted microscope equipped with an A1Rsi confocal scan head and lasers (405, 488, 561, 640 nm) with 20x (NA-0.75, WD 1.0), 40x (NA-0.6, WD 0.14 mm), and 60x (NA-1.27, WD 0.27mm) objectives. Image acquisition and maximum intensity projections were performed using Nikon NIS-Elements v. 5.1 software. Note: the large size of the cells precludes true super resolution microscopy. Many of the FDAA experiments were imaged on a Zeiss Axio Observer SD spinning disk confocal with 10x (NA-0.30, WD 3.2), 40x (NA-0.75, WD 0.71), 63X (NA-1.2, WD 0.28) and 63X (NA-1.4, WD 0.19) objectives with a 488 nm laser. Images were captured using Zen 2 2.0 software. Adjustments to brightness and contrast of confocal stacks was performed in Fiji [125] and of slices/stills in Photoshop. 3D rendering, surface reconstruction, and movie generation for Video 3 were done with Imaris 9.5 (Bitplane). To confirm the types of observations in cellular structure observed during FDAA labeling, additional cells were labeled with WGA with an Alexa Fluor™ 488 conjugate (Life Technologies) and DAPI.

### Co-staining of thin sections for vesicle constituents

Approximately 100 *Ca*. Thiomargarita spp. chains were pooled and fixed with 2% paraformaldehyde in cold PBS with 6.8% sucrose overnight. The cells were dehydrated using a very slow dehydration series with cold ethanol, and the final step using pure ethanol was repeated in triplicate. Then the dehydrated cells were embedded in Technovit® 8100 (Kulzer Technique) and sectioned on a Leica EM UC6 Ultramicrotome producing slices 5 µm in thickness. Individual slices were subjected to different staining procedures. The outer membrane was stained first using polymyxin B labeled with horseradish peroxidase, a 1:50 dilution and incubated overnight [126]. The slides were then wash in PBS wash buffer for 15 minutes and incubated with Alexa Fluor™ 488 Tyramide at 1:100 dilution for 1 hour. Then the slides were washed with both a PBS wash, and H2O wash before counterstaining with FM 4-64 for all lipids. Alternatively, thin sections were stained with DAPI. Image analyses as above on an Olympus BX61 equipped with a UPlanFL N - 100X objective - 1.30 NA Oil Ph3 running CellSens V1.18.

### Transmission electron microscopy method #1

Since, *Ca*. Thiomargarita spp. are essentially a hollow aqueous vacuole for much of the total cell volume, we chose to perform longer fixation steps than typically employed for bacteria. Pooled chains of *Ca*. Thiomargarita spp. cells were fixed in 2% paraformaldehyde in 3.5% NaCl for 24 hours and then were suspended in 2.5% glutaraldehyde, 1M HEPES (4-(2-hydroxyethyl)-1-piperazineethanesulfonic acid) and 3.5% NaCl for two days. Then the cells were washed in Instant Ocean*®* three times and were then resuspended in 2% OsO4 and 3.5% NaCl for two hours. Cells were then washed in HEPES and suspended in 1% uranyl acetate in 3.5% NaCl for one hour followed by another wash series. Cells were then dehydrated through a graded ethanol series (25%, 50%, 75%, 95% and 100% x 3) and were resuspended in a 50:50 mixture of LR White Resin and anhydrous ethanol at 4°C.

After an overnight incubation, the cells were then resuspended in 100% LR White Resin for one hour prior to being placed in gelatin capsules filled with LR White Resin and incubated for one hour at 60°C for polymerization. 40 nm thin sections generated by a Reichert UltraCut S Ultramicrotome. The sections were post-stained with 2% uranyl acetate prior to imaging on a Tecnai™ G2 Spirit BioTWIN transmission electron microscope at an operating voltage of 120 kV. Images were acquired with an Eagle 4 megapixel CCD camera.

### Transmission electron microscopy method #2

Chains of *Ca*. Thiomargarita spp. were transferred three times into petri dishes containing filter-sterilized Instant Ocean, pH 7.8, to remove external debris. Then the chains were pooled and transferred into a 1.5 mL centrifuge tube containing 0.1M cacodylate buffer (pH: 7.4), 2.5% glutaraldehyde, and 2% paraformaldehyde with 5% dimethyl sulfoxide (DMSO) for 15 minutes. The chains were subsequently transferred to fresh buffer and fixative three additional times with 15 minutes intervals to remove residual salts and then were stored at 4°C for 3 hours. The pooled chains were then transferred to the same fixative solution but without DMSO for two weeks and then were shipped to Ohio State University Campus Microscopy & Imaging Facility for processing and imaging. Individual chains of *Ca*. Thiomargarita spp. were transferred to an aqueous room temperature solution of 1% osmium tetroxide and were placed in a Pelco Biowave microwave tissue processing system under vacuum (20” Hg). Each chain was microwaved for two minutes three times with two minutes of rest in-between and then were cooled in an ice bath if the temperature appeared above 27°C. The microwave processing and cooling was repeated two additional times but with only two rounds of microwaving in each series. Then the chains of cells were dehydrated in a graded ethanol series within the microwave at 150 W power. Room temperature 50%, 70%, and 90% ethanol solutions were prepared in sodium acetate to achieve an osmolarity equivalent to seawater (1.16 osmole/L) and an ∼100% ethanol solution was prepared in sodium acetate at an osmolarity of 0.1 osmole/L. Microwave times of 40 seconds each were employed for each increasing concentration of ethanol with the final dehydration step of ∼100% ethanol repeated a second time. The chains were infiltrated with freshly made low viscosity Spurr’s resin in the microwave under vacuum at 200 W. In 5 minutes rounds, the chains were infiltrated with a 1:1 solution of acetone:resin, then by 100% resin for two consecutive rounds followed by cooling at room temperature before a final microwave step with 100% resin. The samples were kept under vacuum for 72 hours and then were transferred to a BEEM capsule containing 100% Spurr’s resin for polymerization in an oven at 65°C for 24 hours. Ultrathin sections were cut by a Leica EM UC6 ultramicrotome and collected on a copper grid. Reynold’s lead citrate and 2% uranyl acetate was utilized for post en bloc staining. Images were acquired with an FEI Technai G2 Spirit transmission electron microscope (FEI), and Macrofire (Optronics) digital camera and AMT image capture software.

### Metagenomics and genome analyses

Two *Ca.* T. nelsonii chains from marine station 23020 sediments stored at 4°C were individually rinsed in sterile seawater and then transferred to UV-treated PBS. TruePrime™ Single Cell Whole Genome Amplification kit (Sygnis) [127] was used to amplify the genomes of *Ca*. Thiomargarita spp. and their microbiomes. Both samples were incubated in 1,4-dithiothreitol for 5 minutes at 25°C followed by 1 minutes at 95°C and then the manufacturer’s instructions were followed. All of the reagents, disposables and equipment were cleaned with DNA-OFF™ (Takara Bio, Inc.) and/or UV sterilized as appropriate [128] prior to amplification. Illumina DNA sequencing libraries were generated using a TruSeq DNA Nano library kit and then the samples were multiplexed with 13 other samples on four lanes of a HiSeq 2500 high output flow cells running v4 chemistry. The resulting number of raw 125x2 reads (550 bps insert) for Sample 1 (aka. ENDO3) was 17,854,737 total paired end reads and for Sample 2 (aka. ENDO5) was 40,206,931 total paired end reads. Residual adapters and phiX were removed with BBDuk version 36.64 [129]. Khmer version 2.0+713.g54c7de6 [130] was then employed to remove low abundance k-mers and PrinSeq-lite 0.20.4 [131] removed low complexity reads and duplicate reads. Both metagenomes were assembled with (meta-)SPAdes-3.10.0 using the k-mers 21,33,55,77 [132]. Genome binning was performed using MyCC using penta-nucleotides and palindromes of hexa-nucleotides [133] on contigs greater than 2000 bps. Complete metagenome assemblies and *Ca*. T. nelsonii bins were submitted to IMG/ER [134] for annotation. Genes of interest were queried against the National Center for Biotechnology Information non-redundant protein sequences database via Blastp with default setting to search for potential associated homologues and examined conserved domain gene regions.

## Data availability

Both the metagenomes (Ga0216254, Ga0216256) and *Ca*. T. nelsonii bins (Ga0309624, Ga0259525) are available at IMG/M. The fastq files are available at https://conservancy.umn.edu/handle/11299/208858.

## Acknowledgements

We would like to thank contributions made by the late Dr. Chibo Chikwililwa, Professor Kurt Hanselmann, Dr. Deon C. Louw, Richard Horaeb, Bronwen Currie, the crew of the R/V Mirabalis, the National Marine Information and Research Centre of the Namibia Ministry of Fisheries and Marine Resources, and the staff and students of Regional Graduate Networks in Oceanography (RGNO) Discovery Camp, and the Marine and Coastal Resources Research Center at the University of Namibia. Also Dr. Jeff Gralnick, Ruth Lee and Dr. Yves Brun for their thoughtful insights. This work was made possible by the skilled staff and resources of the University of Minnesota’s Imaging Center, the UMN Characterization Facility and Genomics Center as well as Ohio State University Center for Electron Microscopy and Analysis and the Campus Microscopy and Imaging Facility. In particular, we thank the formidable research staff Dr. Guillermo Marques, Dr. Jeffrey R. Tonniges and Dr. Giovanna Grandinetti who had to think out of the box to work with very difficult samples. This work was funded by National Science Foundation (1935351) awarded to J.V.B, B.E.F and R.H. and the Simons Foundation Early Career Investigator award (341838), an Alfred P. Sloan Foundation Research Fellowship (BR2014-048) and a McKnight Land Grant Fellowship all awarded to J.V.B. NIH Grant (GM113172) supported the development of the FDAAs. Field sampling was made possible by the RGNO Discovery Camp, which is funded by the Scientific Committee for Oceanographic Research and the Agouron Institute.

## Contributions

B.E.F., J.V.B. and D.J.L. contributed to experimental design. J.V.B., and D.J.L, acquired the samples. M.V.N. provided FDAAs and advice on how to use them. B.E.F., J.V.B., D.J.L. and N.D. performed the FDAA experiments. B.E.F. and J.V.B. conducted the WGA experiments. B.E.F., D.J.L. and J.V.B performed thin section experiments. D.J.L., B.E.F., J.V.B. and R.C.H. performed preparation of TEM samples. R.C.H., B.E.F., and J.V.B. performed TEM analyses. N.D. performed the whole genome amplification. Bioinformatics and genome analysis were by B.E.F with input from B.M. B.E.F. wrote the manuscript with input from all authors.

## Supporting information

**S1 Fig.**
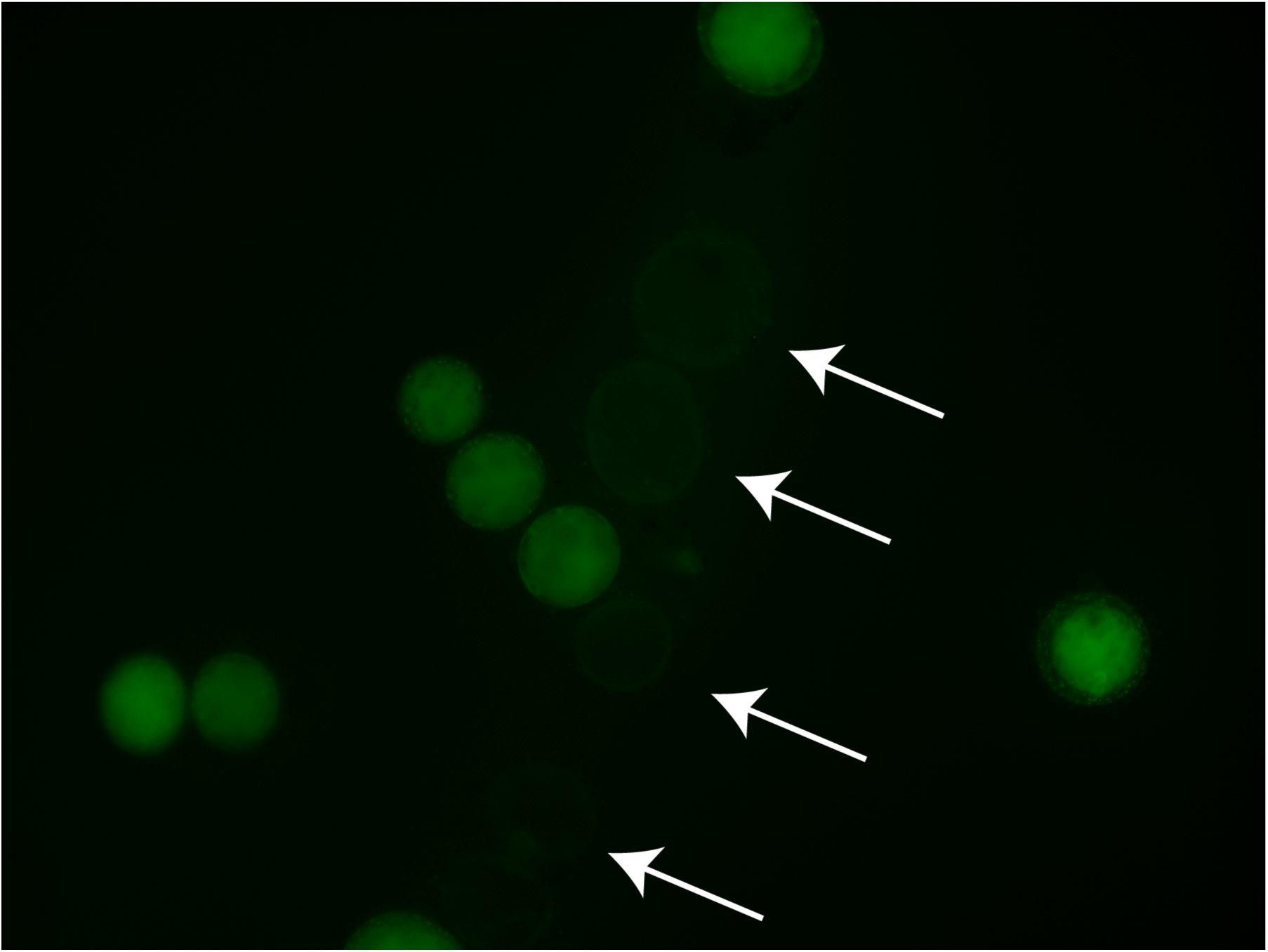
Evidence that FDAAs do not nonspecifically bind cellular material. *Ca.* Thiomargarita sp. Incubated for 1 week with FDAAs. Dead cells, arrows, do not incorporate the FDAAs.

**S2 Fig.**
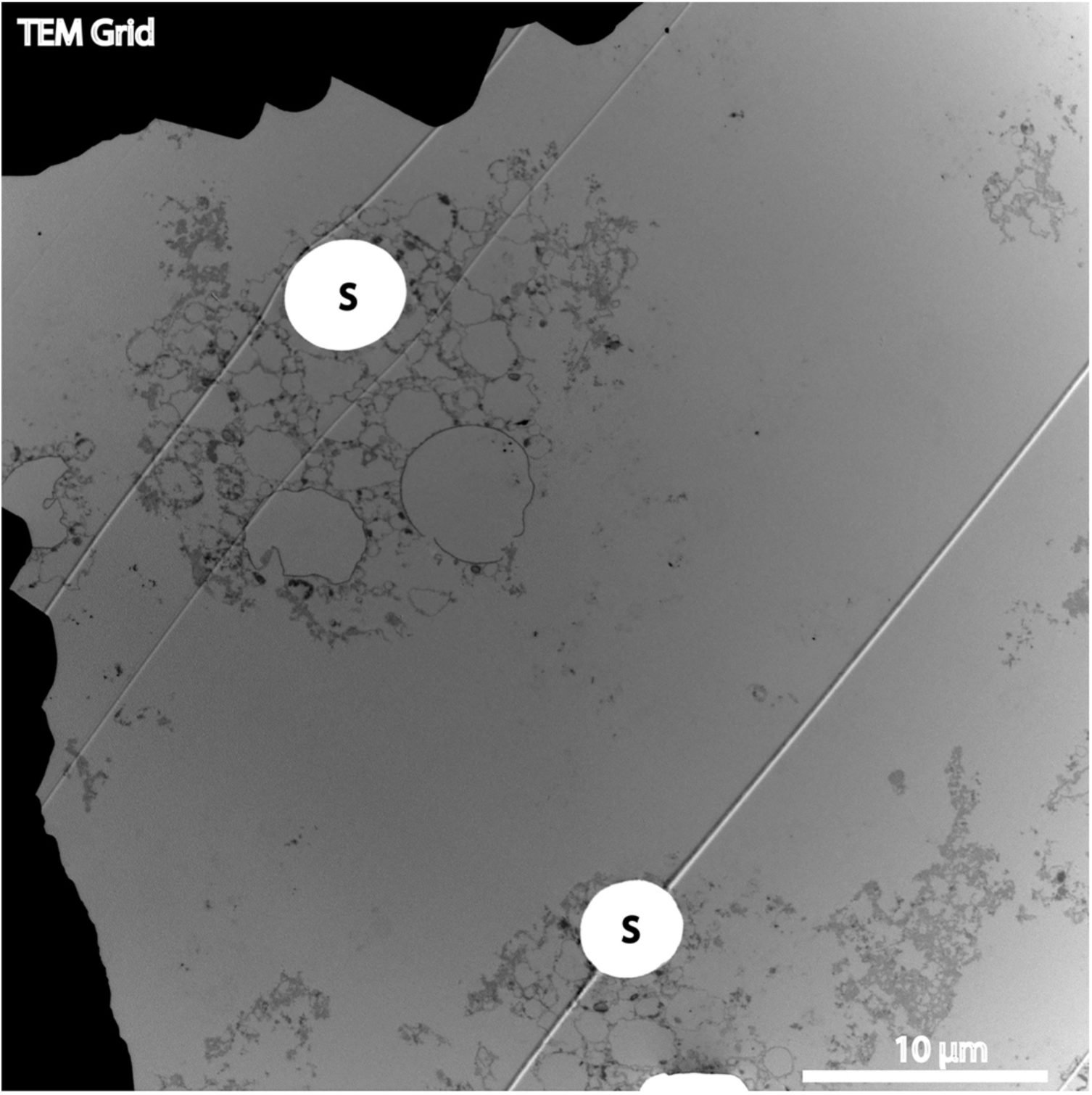
Transmission electron microscope micrograph of clusters of VLFs in *Ca*. Thiomargarita spp.. As depicted here, some clusters of VLFs can become detached from the cellular envelope. Some large VLFs contain sulfur (S) but many do not. Some deformation of vesicles, particular large vesicles, occurred during the infiltration of resin. Some dark matter, like DNA and RNA, are within vesicles but some were not. Note, the lines across the micrograph are from the imperfections in the sectioning blade.

**S3 Fig.**
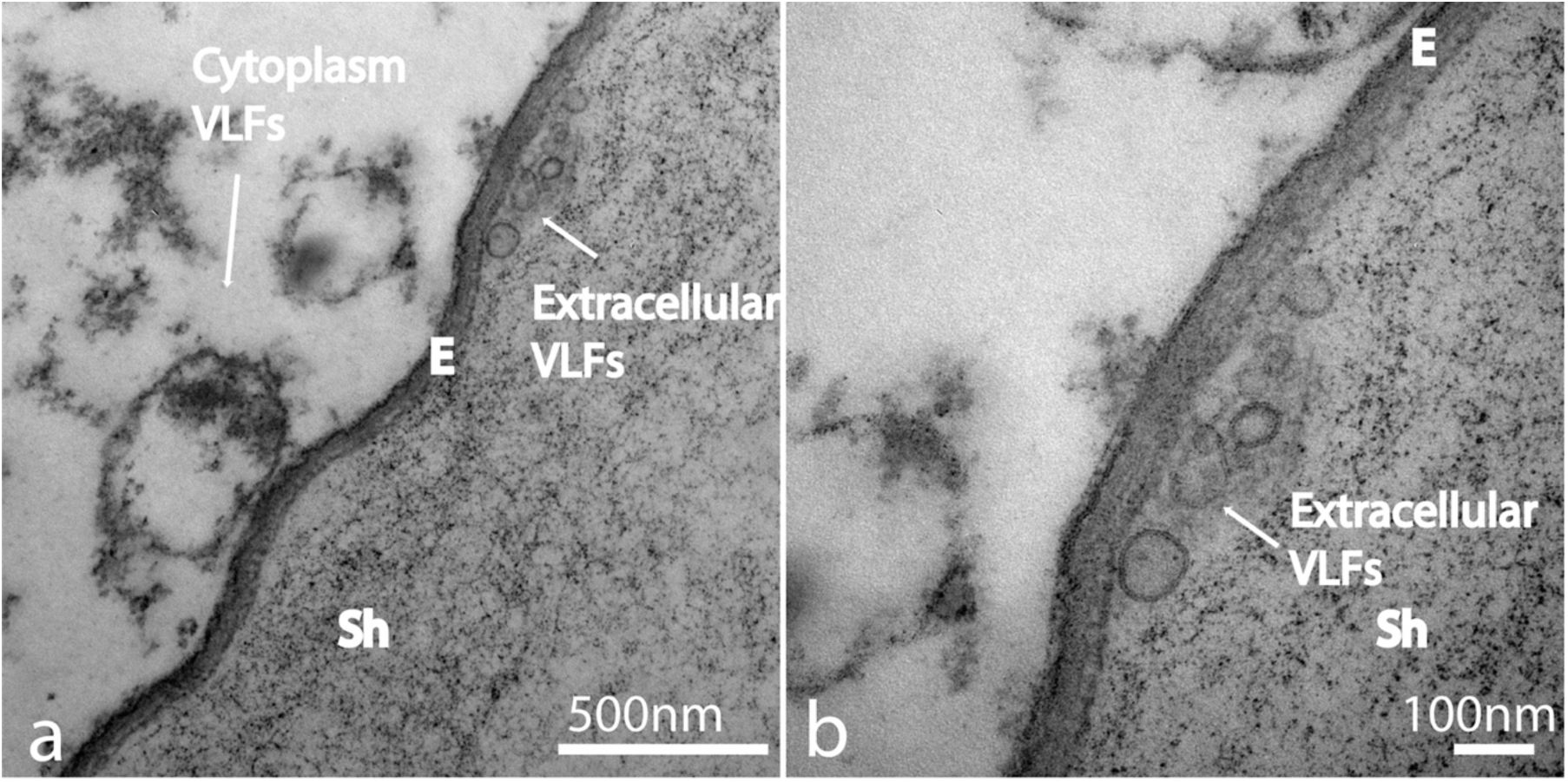
Transmission electron microscopy images of the cellular envelope of *Ca*. Thiomargarita spp. revealing the production of outer membrane vesicles exterior to the outermost layer (S-layer). (a) Outer membrane vesicles appear smaller in size to intracellular vesicles. (b) Outer membrane vesicles appear to have bi-layered walls. E = envelope and Sh = sheath.

**S1 Movie. Confocal Z-stack of a chain Ca. Thiomargarita spp. incubated with FDAAs for 30 minutes revealed reductive cell division as seen in** **Figure 2c**.

**S2 Movie. Confocal Z-stack of a Ca. Thiomargarita spp. cell undergoing cell division, as seen in** **Fig. 4, reveal PG bound vesicles detached from the envelope.**

**S3 Movie. 3D rendering and surface reconstruction of cell, featured in** **Fig. 4** **and S2 Movie, explores the division plane and intracellular environment.**

**S4 Movie. Confocal Z-stack of a *Ca*. Thiomargarita spp. cell, featured in** **Fig. 5a-b** **stained with wheat germ agglutin.**

**S5 Movie. Confocal Z-stack of a *Ca*. Thiomargarita spp. cell, featured in** **Fig. 5a-b** **and S4 Movie, stained with DAPI.**

**S1 Data. Putative genes in genomes *Ca*. T. nelsonii pertaining to chromosome partitioning, outer membrane, cell division, PG synthesis and degradation and cell elongation.**

**S2 Data. Putative genes in genomes *Ca*. T. nelsonii pertaining to membrane lipids, cytoskeletal proteins, flotillins, small GTPases, as well as other genes queried for in the genomes.**

